# Ephrin-B2 promotes nociceptive plasticity and hyperalgesic priming through EphB2-MNK-eIF4E signaling in both mice and humans

**DOI:** 10.1101/2024.02.21.581414

**Authors:** Eric T. David, Muhammad Saad Yousuf, Hao-Ruei Mei, Ashita Jain, Sharada Krishnagiri, Kolluru D. Srikanth, Gregory Dussor, Matthew B. Dalva, Theodore J. Price

## Abstract

Ephrin-B-EphB signaling promotes pain through signaling between dorsal root ganglion (DRG) neurons and spinal cord neurons in the dorsal horn, and through signaling between peripheral cells and EphB receptors expressed by DRG neurons. Previous findings link ephrin-B expression in painful peripheral tissues in patients to chronic pain, suggesting the clinical significance of this signaling, but the direct effects of ephrins on DRG neurons have not been widely studied. We hypothesized that ephrin-B2 would promote nociceptor plasticity and hyperalgesic priming through MNK-eIF4E signaling, a critical mechanism for nociceptive plasticity induced by growth factors, cytokines and nerve injury. Our work demonstrates that ephrin-B2-EphB2 signaling drives activation of MNK-eIF4E in DRG neurons to cause an enhanced response to inflammatory mediator signaling in both mice and humans and hyperalgesic priming in two models in mice. Both male and female mice developed dose-dependent mechanical hypersensitivity in response to ephrin-B2, and both sexes showed hyperalgesic priming when challenged with PGE_2_ injection into the same hindpaw. Acute nociceptive behaviors and hyperalgesic priming were blocked in mice lacking MNK1 (*Mknk1* knockout mice) and by the MNK inhibitor eFT508. Similar effects on hyperalgesic priming were seen in a dural injection model. We generated a sensory neuron specific knockout of EphB2 using Pirt-Cre mice and found that these mice lacked responses to ephrin-B2 injection. We used Ca^2+^-imaging to determine direct effects of ephrin-B2 on DRG neurons and found that ephrin-B2 treatment enhanced Ca^2+^ transients in response to PGE_2_ which were absent in DRG neurons from MNK1^−/−^ and EphB2-Pirt^Cre^ mice. In experiments on human DRG neurons we found that ephrin-B2 increased eIF4E phosphorylation and enhanced Ca^2+^ responses to PGE_2_ treatment, both of which were blocked by eFT508 treatment. We conclude that ephrin-B2 acts directly on mouse and human sensory neurons to induce nociceptor plasticity via MNK-eIF4E signaling. The findings offer insight into how ephrin-B signaling promotes pain, and suggests treatment avenues for prevention or reversal of chronic pain associated with EphB activation in sensory neurons.

**Graphical Abstract:** 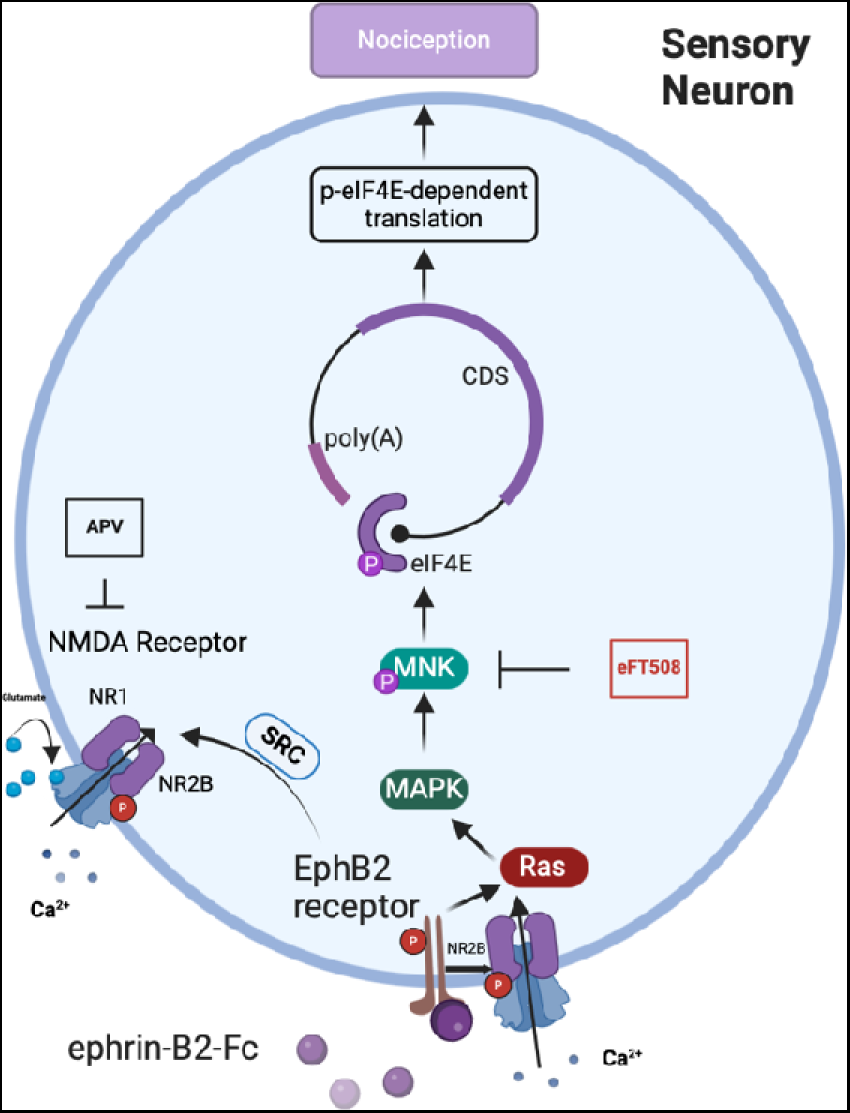

## Introduction

EphB class of receptor tyrosine kinases subclass B (EphB) and their ligands, the Ephrin-Bs, are key signaling proteins in neuronal development, synaptogenesis and synaptic plasticity [26] that have been implicated in nervous system pathologies such as neuropathic and post-injury pain [1; 10; 17; 24; 30; 37; 66; 70; 79]. Ephrin-Bs and EphBs signal by binding in transcellular multimeric complexes to induce bidirectional signaling in the cell expressing the ephrin-B ligand and the EphB receptor [26]. In the nervous system, a key consequence of ephrin-B-EphB signaling occurs through EphB1 and EphB2 coupling to the NMDA receptor (NMDAR) at postsynaptic sites [19; 24; 52; 74]. It is now well-recognized that ephrin-B-EphB signaling produces NMDAR-dependent nociceptive signal amplification at spinal cord dorsal horn excitatory synapses [10; 24; 30; 70; 79] and at medullary dorsal horn synapses receiving inputs from trigeminal (TG) afferents that are important for headache pain [72]. While this central nervous system mechanism has been described, the role of ephrin-B-EphB signaling in the peripheral nervous system is not as extensively studied. Ephrin-B ligands are expressed by DRG neurons, including nociceptors, and they increase after injury in rodent DRG neurons [30; 66; 79]. They are also found in other peripheral cell types [68], and their expression can be increased in painful diseases in humans, like pancreatic cancer [73] and rheumatoid and osteoarthritis [31]. Moreover, increased *EFNB2* gene expression is linked to pain in treatment resistant rheumatoid arthritis [8]. Both EphB1 and EphB2 are expressed in mouse DRG neurons, and similar expression is observed in human DRG, but *EPHB2* mRNA expression is higher in both mice and humans [12; 69; 78]. Both receptors are also expressed in the human TG, suggesting a peripheral role in TG-driven pain, like headache [75]. Only one previous study has extensively studied ephrin-B2-EphB signaling in rodent DRG neurons finding evidence for src and extracellular signaling regulated kinase (ERK) kinase engagement and increased NMDAR signaling [37]. We sought to better understand how ephrin-B2 might act on both mouse and human DRG neurons with the hypothesis that activating this receptor would lead to mitogen activated protein kinase (MAPK) signaling-dependent sensitization of DRG neurons via the EphB2 receptor.

Canonical EphB signaling involves tyrosine phosphorylation events and biochemical interactions at specific domains of the protein on both the intracellular and extracellular side of the receptor [26]. However, other signaling mechanisms can be engaged by ephrin-B-EphB binding, including MAPK signaling [11; 35; 54; 62]. The MAPK-ERK and its downstream effector for regulation of mRNA translation, mitogen activated kinase interacting kinase (MNK1/2), have been extensively implicated in nociceptive plasticity after injury, and the control of sensory neuron excitability [18; 28; 34; 37; 38; 41; 44; 48; 49]. MNK1/2 phosphorylate an RNA binding protein called eIF4E that binds to the 5’ cap structure of mRNAs to regulate the translation of a subset of mRNAs that are involved in inflammation and nociceptor excitability. Because ephrin-B upregulation is associated with painful disease in humans [8; 31; 73], we reasoned that MNK1/2 activation and eIF4E phosphorylation downstream of EphB activation could be a key mechanism driving nociceptive plasticity in response to experimental treatment with ephrin-B2. We tested this hypothesis using mouse models of hyperalgesic priming, which assess behavioral manifestations of changes in signaling and gene expression in sensory neurons [55; 58; 59; 71], and model some aspects of the transition from acute to chronic pain [55; 59]. We also used Ca^2+^ imaging studies on mouse and human DRG neurons to understand direct effects on these neurons and to provide direct clinical translational value to our experiments [60]. Our findings provide clear evidence that ephrin-B2 promotes nociceptive plasticity via EphB2 expressed by sensory neurons and via MNK1/2 signaling in both mice and humans. The work substantiates that manipulating EphrinB-EphB signaling in the peripheral nervous system can be a target mechanism for pain treatment.

## Methods and Materials

### Animals

All animal procedures were approved by the Institutional Animal Care and Use Committee at the University of Texas at Dallas (protocol number 14-04). Seven- to 12-week-old male and female mice were used in this study. Animals were either bred in the Animal Research Facility at the University of Texas at Dallas or through Charles River laboratories. Strains used in experiments include C57BL/6J, MNK^−/−^ on a C57BL/6J background, and EphB2-Pirt^+/-^ on a C57BL/6J background. To generate EphB2^flox^ mice, loxP sites were inserted flanking the *Ephb2* [67] gene and crossed with Pirt^Cre^ [29] on a C57BL6/J background to make sensory neuron specific knockout mice for EphB2. Animals were housed in same-sex groups of 4 on a 12 hour light/dark cycle, with food and water available *ad libitum*.

### Organ donors

All human tissue procurement procedures were approved by the Institutional Review Board at the University of Texas at Dallas. Human DRGs were recovered from organ donors through collaboration with Southwest Transplant Alliance. Upon recovery, which has been described previously [64; 69], the DRGs were immediately kept in N-methyl-D-glucamine supplemented artificial cerebral spinal fluid (aCSF) on wet ice as previously described [69] until enzymatic digestion for generation of cultures for Ca^2+^ imaging.

### Drugs

Recombinant mouse ephrin-B2-Fc (496-EB-200), recombinant human ephrin-B2 (7397-EB-050), and human IgG (110-HG-100), were purchased from R&D systems. Prostaglandin E_2_ (PGE_2_) was obtained from Cayman Chemical (14010). eFT508 (HY-100022, MedChem Express), was reconstituted as previously described [63]. Stock ephrin-B2-Fc was reconstituted with sterile 0.9% saline and diluted in sterile 0.9% saline to either 1 ng, 10 ng, or 100 ng dose per injection and did not undergo multiple freeze thaw cycles. Intraplantar injections were given as previously described [4]. PGE_2_ was reconstituted as previously described [4]. PGE_2_ was diluted into sterile 0.9% saline to a final dose of 100 ng per injection [4]. Recombinant human ephrin-B2 was reconstituted in sterile 0.9% saline. Prior to pre-treatment, ephrin-B2 was conjugated to human IgG for 30 minutes prior to incubation as previously described [24]. For calcium imaging, stock PGE_2_ was diluted in normal recording solution (described below) to a final concentration of 50 nM. All drugs used for *in vivo* use were sterile filtered with a 0.22µm filter and kept on ice until immediately before injection.

### von Frey testing

Animals were habituated in an acrylic box with mesh flooring for 1 hour before testing. The paw withdrawal threshold was determined using the up-down method with calibrated von Frey filaments (37450-275, Ugo Basile) [16]. Baseline paw withdrawal thresholds were determined on the left hindpaw prior to any treatments. The experimenter was blinded to experimental conditions.

### Heat hyperalgesia testing

Thermal heat sensitivity was determined using the Hargreaves method (IITC Life Science) [25]. Mice were placed on 29° C glass for 30 minutes before each test using the IITC Model 390 plantar test apparatus. A focused high-intensity light was aimed at plantar surface of the left hind paw of mice. Light intensity was set to 35% of maximum with a cutoff of 20 seconds. Hindpaw withdrawal latency was measured prior to treatment, and at 1, 3, 5, 24, and 72 hours after treatment with three tests averaged for each timepoint.

### Supradural injections

Male and female mice at ages 4-6 weeks were habituated prior to experimentation. Injections of ephrin-B2-Fc were given to the dura as previously described [15]. Mice were tested using calibrated von Frey filaments using the up-down method [16]. Mice were anesthetized with isoflurane for less than 2 min. The injection was performed through a modified internal cannula (P1 Technologies, Roanoke, VA, USA, part #8IC313ISPCXC, Internal Cannula, standard, 28 gauge, fit to 0.5 mm) and the total volume of injection was 5 μL. The injector projection length, measured by digital calipers, ranged from 0.5 mm to 0.6 mm, enabling us to reach the dura mater through the intersection of the lambdoidal and sagittal. After baseline was established, animals were tested after injection at 1, 3, 5, 24, and 72 hours. Once mice returned to baseline, a subsequent injection of PGE_2_ was given onto the dura and mechanical testing was done at 1, 3, and 24 hours.

### Hyperalgesic priming

To establish hyperalgesic priming, 100 ng of ephrin-B2-Fc was injected intraplantarly into the left hindpaw. Paw withdrawal thresholds were subsequently measured at 1, 3, 5, 24, and 72 hours. Mice were then injected intraplantarly with 100ng of PGE_2_ and then tested for mechanical paw withdrawal threshold at 1 and 3 hours after injection.

### Primary Neuronal Cultures

Mouse DRGs were dissected from adult male and female mice at 4-6 weeks of age bred at the University of Texas at Dallas Animal Research Facility. DRGs were suspended in Hanks balanced salt solution (HBSS) without calcium and magnesium (14170161) before culturing. DRGs were incubated at 37° C for 25 minutes in 1 mg/mL papain (10108014001, Sigma-Aldrich), followed by incubation at 37° C for 25 minutes in 3 mg/mL collagenase type 2 (LS004174, Worthington Biochemical) and 2 mg/mL Dispase II (4942078001, Sigma-Aldrich). DRGs were then triturated in DMEM/F12 plus GlutaMAX (11-320-033, Fisher Scientific) and passed through a 70 μm cell strainer and resuspended. Cells were plated onto either 1mg/ml Poly-D-Lyisine coated coverslips or 35 mm culture dish with 14 mm glass microwell. Dissociated neurons were plated and allowed to adhere for 2 hours at 37°C in 5% CO_2_ wells were flooded with DMEM/F12 plus GlutaMAX culture media containing 1% penicillin/streptomycin (15070063, Thermofisher), 10 ng/mL mouse nerve growth factor (01-125, EMD Millipore, NGF), and 3 μg/mL 5-fluoro-2′-deoxyuridine + 7 μg/mL uridine (FRDU, F0503, Sigma-Aldrich). Cultured cells were then kept at 37°C and 5% CO_2_ in an incubator with culture media changes at 48 hours with supplemented media and NGF and FRDU until further experimentation.

Human DRGs were prepared and cultured as previously reported [76]. In brief, human DRGs were microdissected with scissors into 1-2 cm pieces while immersed in NMDG-aCSF. The DRG pieces were incubated in 5 ml HBSS containing 2 mg/mL Stemzyme (LS004106, Worthington Biochemical) and DNAse I (LS001239, Worthington Biochemical) at 4 µg/mL at 37° C in a shaking water bath for up to 12 hours. Cells were then passed through a 100 μm cell strainer and pipetted onto a solution of 10% sterile filtered bovine serum albumin (BSA, BioPharm) in HBSS. Cells were centrifuged for 5 min at 900xg then resuspended in BrainPhys media (05790, STEMCell Technologies) with 1% penicillin-streptomycin (15070063, ThermoFisher), 2% SM1 (05711, STEMCell Technologies), 1% N2 (07152, STEMCell Technologies), and 1% GlutaMax (Gibco). A 50 µl cell droplet was incubated for 2 hours at 37° C. Wells were flooded with BrainPhys media with supplements and 10 ng/mL human NGF (256-GF-100, R&D Systems), and 3 μg/mL FRDU and incubated at 37° C and 5% CO_2_ until further experimentation.

### Calcium Imaging and Analysis

Mouse cultures prepared as described above were grown on 35mm glass coverslips coated with 1 mg/mL Poly-D-Lysine (P0899, Sigma Aldrich) and imaged at 48 hours after plating. Cells were loaded with Fura-2 AM at 1 μg/mL (108964-32-5, Life Technologies) in HBSS or Fluo-4 AM at 10 μM in (ThermoFisher) 2% Pluronic F-127 (ThermoFisher) for 1 hour prior to imaging. During pre-treatment, cultures were treated with either pre-clustered ephrin-B2, IgG Control or pre-clustered ephrin-B2 with eFT508 prior to loading dye and kept at 37°C and 5% CO_2_. After incubation, Fura-2 AM or Fluo-4 AM was replaced with normal recording solution (135 mM NaCl, 5 mM KCl, 10 mM HEPES, 2 mM CaCl_2_, 1 mM MgCl_2_ and 10 mM glucose osmolarity adjusted to 300 ± 5 mOsm and pH adjusted to 7.4 with N-methyl-D-glucamine) for 20 minutes prior to imaging. In human cultures, recording solution (121 mM NaCl, 4.2 mM KCl, 1.1 mM CaCl_2_, 1 mM MgSO_4_, 29 mM NaHCO_3_, 0.45 mM NaH_2_PO_4_-H_2_O, 0.5 mMNaH_2_PO_4_-7H_2_O, 20 mM Glucose, 10 mM HEPES) was bubbled in carbogen for 30 minutes prior to recording (modified from [9]). High K^+^ solution was made by adjusting KCl to 50 mM and subtracting the difference from the NaCl concentration to maintain osmolarity. The recordings were performed using an Olympus IX73 inverted microscope at 20X magnification and the Meta-Fluor Fluorescence Ratio Imaging Software. Cells that exhibited 10% Fura-2-AM ratiometric change (340/380nm) or intracellular Fluo-4 AM in Ca^2+^ levels in response to 50 mM K^+^ treatment were considered as responsive. All high K^+^ or PGE_2_ responding neurons were identified if a 10% change in Ca^2+^ levels occurred.

### Immunocytochemistry

Human cultures were prepared as described above and plated onto 12 mm glass coverslips for ≥48 hours and up to 96 hours with normal media change at 48 hours. Cells were fixed with 10% formalin for 15 minutes and washed three times with 1X PBS. Cells were blocked with 10% normal goat serum (NGS) in 0.1% Triton X dissolved in 1X PBS. Cells were incubated overnight at 4° C in primary anti-peripherin raised in chicken (1:1000, EnCor Biotechnology #cpca-peri) and anti-p-eIF4E raised in rabbit (1:1000, Fisher Scientific #ab76256). Cells were washed three times with 1X PBS for 5 minutes each. Secondary antibodies DAPI (1:5000, Cayman Chemical #14285), Alex Fluor 488 (1:2000, Invitrogen #A-11039) and Alex Fluor 555 (1:2000, ThermoFisher #A21428) were incubated at room temperature for 2 hours. Cells were washed three times with 1X PBS prior to mounting on uncharged slides with ProLong Gold Antifade mounting medium (Fisher Scientific #P36930) and kept from light until imaging.

### Confocal Microscopy imaging and analysis

Images were captured using an Olympus FV3000 confocal microscope. Filters were selected for detection of Alexa Fluor 405, Alexa Fluor 488, and Alexa Fluor 555 using multitrack scanning. Images were taken with a 20X objective lens. All setting parameters throughout imaging were kept consistent. A minimum of 3 images were taken per well. Raw images were analyzed using Olympus CellSens software and manually assessed for expression of p-eIF4E. Peripherin signal was used to verify neuronal cells. Regions of interest of each neuron were drawn using the ellipse tool, and p-eIF4E was quantified using mean grey intensity value. Background levels from negative control were used to subtract before analysis. Levels of p-eIF4E were compared between treatment groups.

### Statistics

All statistical analysis was performed using GraphPad Prism version 9.4.1 and all data are shown as mean ± SEM. For behavioral analysis, differences between groups were determined by repeated measures two-way ANOVA with Bonferroni’s post hoc test α = 0.05. For Ca^2+^ imaging experiments, chi-squared analysis was performed between treatments. Latency to peak responses and immunocytochemistry data were determined using one-way or two-way ANOVA with Bonferroni’s post hoc test α = 0.05. Chi-squared statistic is given in the results section. All other statistical tests are indicated in figure panels.

## Results

### Ephrin-B2 induces acute mechanical hypersensitivity and hyperalgesic priming in both male and female mice, but only heat hyperalgesia in males

We first tested the effects of ephrin-B2 injected into the hindpaw to assess mechanical and heat threshold responses and subsequent development of hyperalgesic priming. Animals were given ascending doses of ephrin-B2 at 1, 10, and 100 ng and tested as shown in **Fig 1A**. Following the resolution of initial mechanical hypersensitivity, animals receive an injection of 100 ng PGE_2_ to assess the presence of hyperalgesic priming. Male mice developed acute mechanical hypersensitivity in response to ephrin-B2 injected intraplantarly at 10 and 100 ng (**Fig 1B**). Additionally, males showed mechanical hypersensitivity to PGE_2_ after receiving the 100 ng intraplantar dose of ephrin-B2, but not vehicle or lower doses (**Fig 1B**). To determine if ephrin-B2’s nociceptive inducing effects were sex specific, we also tested female mice at 1, 10, and 100 ng doses and observed similar effects to those seen in males (**Fig 1C**). Again, in females, only the 100 ng group displayed evidence of hyperalgesic priming when challenged with hindpaw injection of 100 ng PGE_2_ (**Fig 1C**). Males tested for heat hyperalgesia only displayed evidence of hyperalgesia in the 100 ng ephrin-B2 dose group (**Fig 1D**). Female mice did not display heat hyperalgesia with any dose of ephrin-B2 (**Fig 1E**). We did not test for heat hyperalgesia in hyperalgesic priming because of this sex difference, and because heat hyperalgesia in response to PGE_2_ has not been reported in similar models [43].

**Figure 1.**
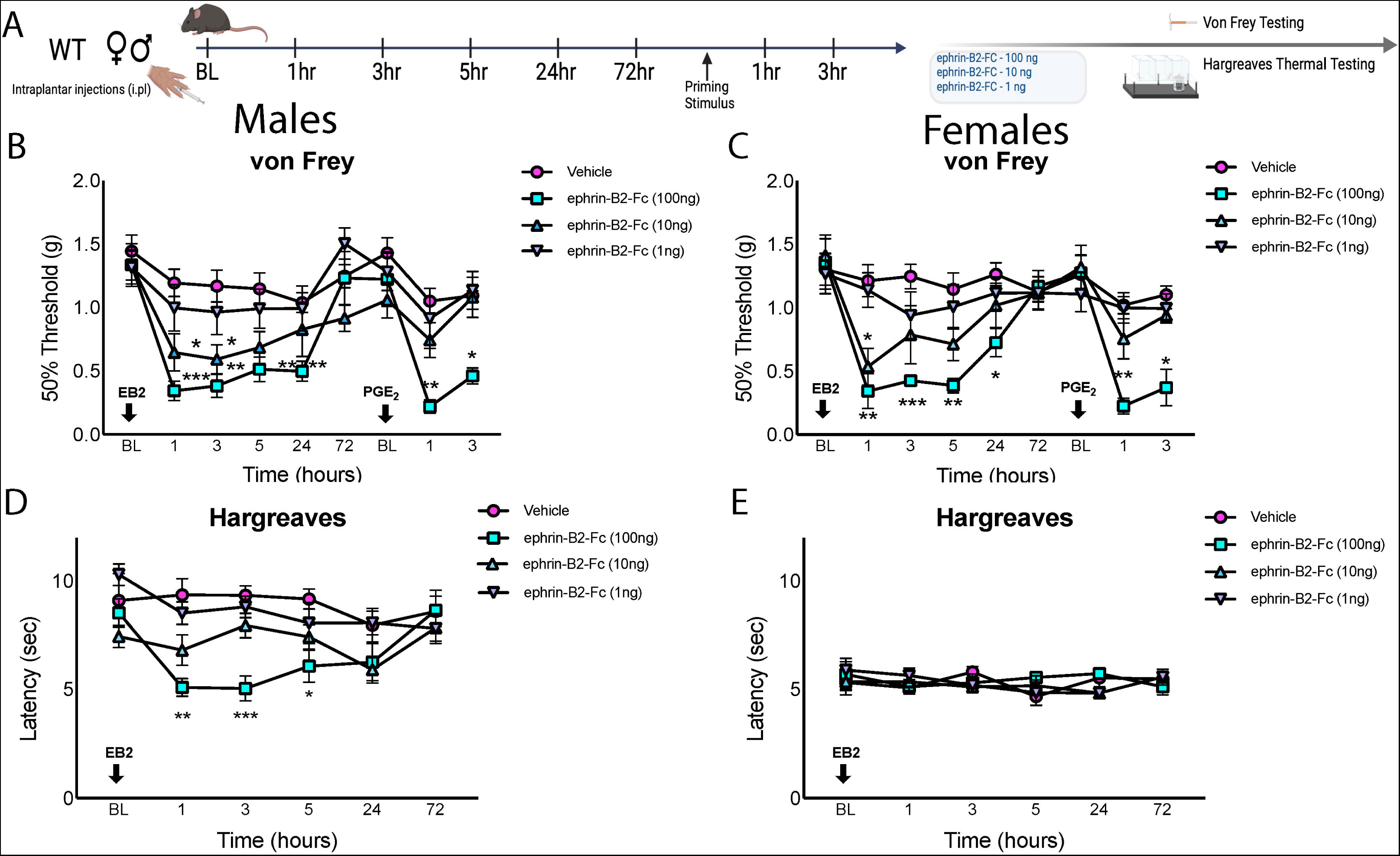
Response to intraplantar injection of ephrin-B2-Fc or vehicle in both male and female mice. A, Overview of behavioral testing procedure. B, Mechanical thresholds after ephrin-B2-Fc intraplantar injection in male mice. Mice received an intraplantar injection of 100 ng of PGE_2_ after initial mechanical hypersensitivity had resolved to assess the presence of hyperalgesic priming. C, Mechanical thresholds after ephrin-B2-Fc intraplantar injection in female mice. Hyperalgesic priming was assessed over the same time course as in male mice. D, Male mice were injected with ephrin-B2-Fc or vehicle and tested for thermal hyperalgesia using the Hargreaves method. E, Female mice were injected with ephrin-B2-Fc or vehicle and tested for thermal hyperalgesia using the Hargreaves method. Males and females n = 6/group, asterisk represent significant differences between groups. EB2 = ephrin-B2-Fc. Repeated measures two-way ANOVA with Bonferroni’s *post hoc* test: *p < 0.05, **p < 0.01, ***p < 0.001.

### Ephrin-B2 induced nociceptive behavioral effects are MNK1-dependent in male and female mice

We then tested if the ephrin-B2-induced mechanical hypersensitivity and heat hyperalgesia in males were driven through MNK-eIF4E signaling. To do this, we injected ephrin-B2 at 100 ng in wild-type and MNK1^−/−^ and assessed the presence of hyperalgesic priming with subsequent 100 ng injection of PGE_2_ We found MNK1^−/−^ males intraplantarly injected with ephrin-B2 do not develop acute mechanical hypersensitivity or hyperalgesic priming (**Fig 2A**). Female MNK1^−/−^ mice also did not develop acute mechanical hypersensitivity or hyperalgesic priming to 100 ng ephrin-B2 intraplantar injection (**Fig 2B**). Male MNK1^−/−^ mice did not display thermal hyperalgesia after receiving 100 ng ephrin-B2 intraplantarly (**Fig 2C**). Female wild-type and MNK1^−/−^ displayed a significant overall effect following ephrin-B2 injection (**Fig 2D**).

**Figure 2.**
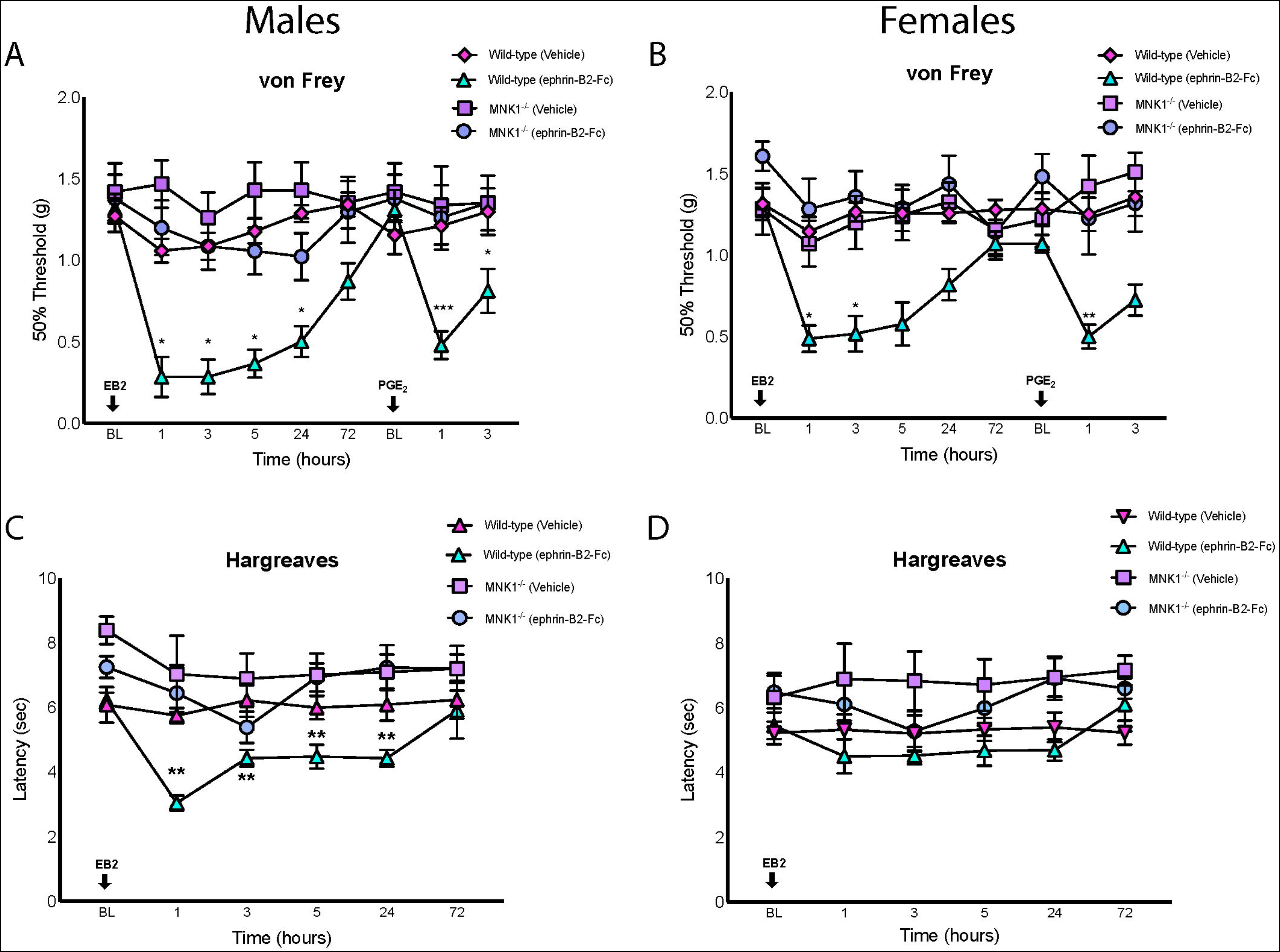
MNK1 is necessary for ephrin-B2-Fc induced mechanical hypersensitivity, thermal hyperalgesia, and hyperalgesic priming. A, B, Wild-type or MNK1^−/−^ male mice were given an intraplantar injection of 100ng ephrin-B2-Fc or vehicle followed by an injection of 100ng PGE_2_ after initial hypersensitivity had resolved. C, D, Wild-type or MNK1^−/−^ mice were given intraplantar injections of either ephrin-B2-Fc or vehicle and tested for thermal hyperalgesia using the Hargreaves method. Stars represent statistical significance. Males and females n = 6/group. EB2 = ephrin-B2-Fc. Repeated measures two-way ANOVA with Bonferroni’s *post hoc* test: *p < 0.05, **p < 0.01, ***p < 0.001

### eFT508 treatment blocks ephrin-B2 induced nociceptive behavioral effects in both sexes

Effects in MNK1^−/−^ mice suggest that acute pharmacological inhibition of MNK should reduce ephrin-B2-driven nociceptive behaviors. To formally test this, we used the known MNK1/2 inhibitor eFT508 with oral dosing [33; 41; 57; 63] prior to ephrin-B2 intraplantar injection. Acute mechanical hypersensitivity and hyperalgesic priming were blocked with eFT508 (10 mg/kg) pre-treatment (**Fig 3A**). Female mice given eFT508 pre-treatment also did not display significant mechanical hypersensitivity or hyperalgesic priming (**Fig 3B**). Male mice given eFT508 prior to ephrin-B2 did not display thermal hyperalgesia, showing that this sex-specific response was also MNK1-mediated (**Fig 3C**). Female mice did no develop thermal hyperalgesia in response to ephrin-B2 (**Fig 3D**).

**Fig 3:**
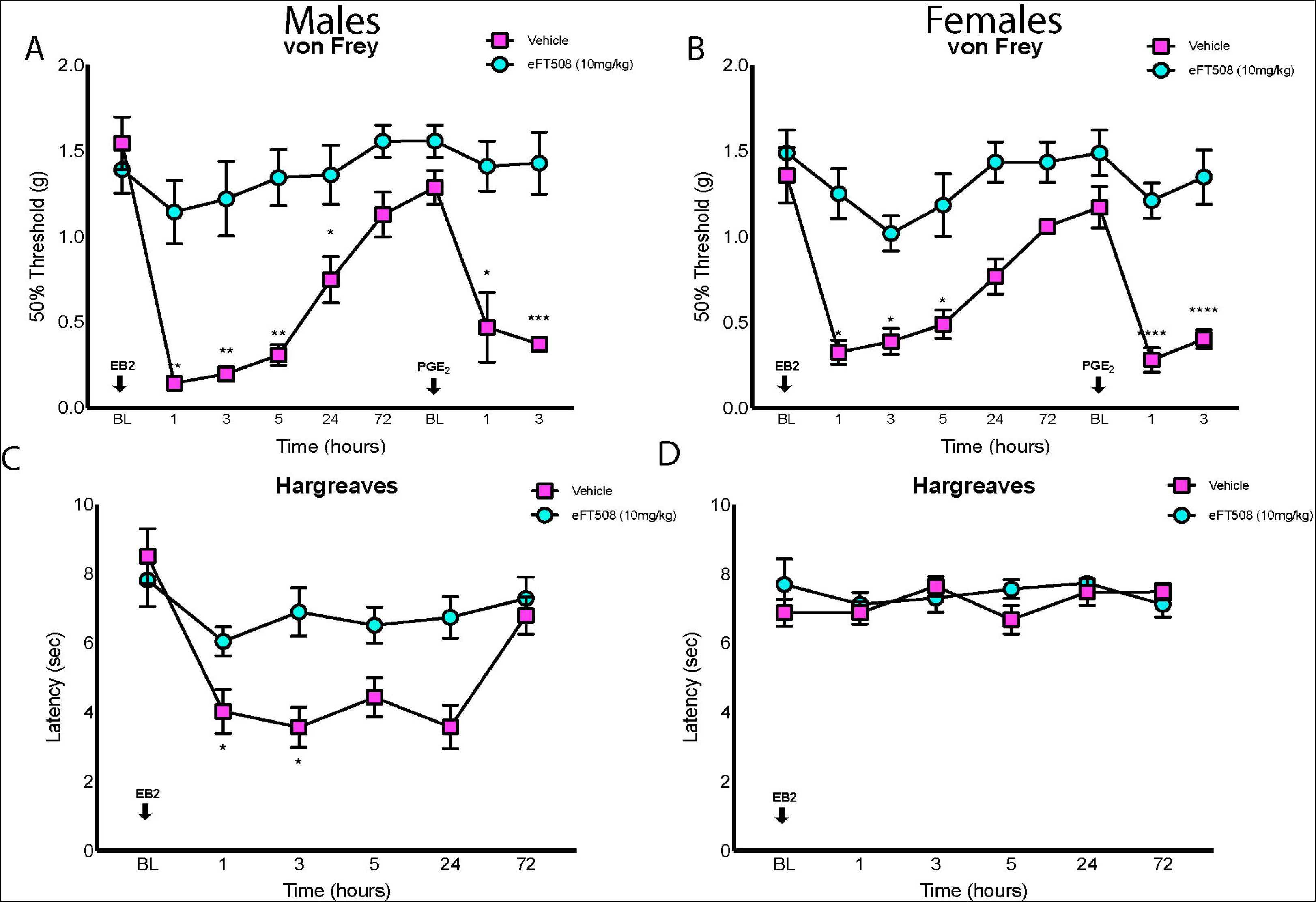
MNK1/2 inhibition with eFT508 blocks ephrin-B2-driven nociceptive behaviors. A, B, Mice given an oral dose of eFT508 (10 mg/kg) compared to vehicle 30 minutes prior to receiving an intraplantar injection of 100 ng of ephrin-B2-Fc show attenuated mechanical hypersensitivity. After initial hypersensitivity was resolved, mice received an intraplantar injection of 100 ng of PGE_2_ to assess the presence of hyperalgesic priming. C, D, Mice were given an oral dose of eFT508 or vehicle 30 minutes prior to receiving an intraplantar injection of ephrin-B2-Fc and tested for thermal hyperalgesia. Stars represent statistical significance. Males and females n = 6/group. EB2, ephrin-B2-Fc. Repeated measures two-way ANOVA with Bonferroni’s *post hoc* test: *p < 0.05, **p < 0.01, ***p < 0.001, ****p < 0.0001.

### Ephrin-B2-induced nociceptive behavioral effects require EphB2 receptor expression on sensory neurons

Previous studies have mostly implicated EphB1 in ephrin-B2-mediated nociceptive effects, but these studies have relied heavily on global knockout of the gene [17; 23; 36] or implicated a spinal action of EphB1 receptors [10]. Unbiased methods demonstrate that both EphB1 and EphB2 are expressed in the DRG [12; 69; 78], motivating us to examine the potential role of sensory neuronal EphB2 in the nociceptive effects of ephrin-B2 treatment. To do this we crossed floxed EphB2 mice with Pirt-Cre mice because Pirt is predominantly expressed by sensory neurons [29]. Male EphB2-Pirt^Cre^ mice injected with ephrin-B2 intraplantarly did not display significant mechanical hypersensitivity or hyperalgesic priming while EphB2-Pirt^+/+^ mice did (**Fig 4A**). Female EphB2-Pirt^Cre^ mice injected with ephrin-B2 also did not develop mechanical hypersensitivity or hyperalgesic priming while EphB2-Pirt^+/+^ showed both, consistent with previous observations (**Fig 4B**). Male EphB2-Pirt^Cre^ mice did not develop significant thermal hyperalgesia while EphB2-Pirt^+/+^ mice did at 1 and 3 hours after treatment (**Fig 4C**). As previously shown, females injected with ephrin-B2 intraplantarly did not develop thermal hyperalgesia (**Fig 4D**).

**Fig 4:**
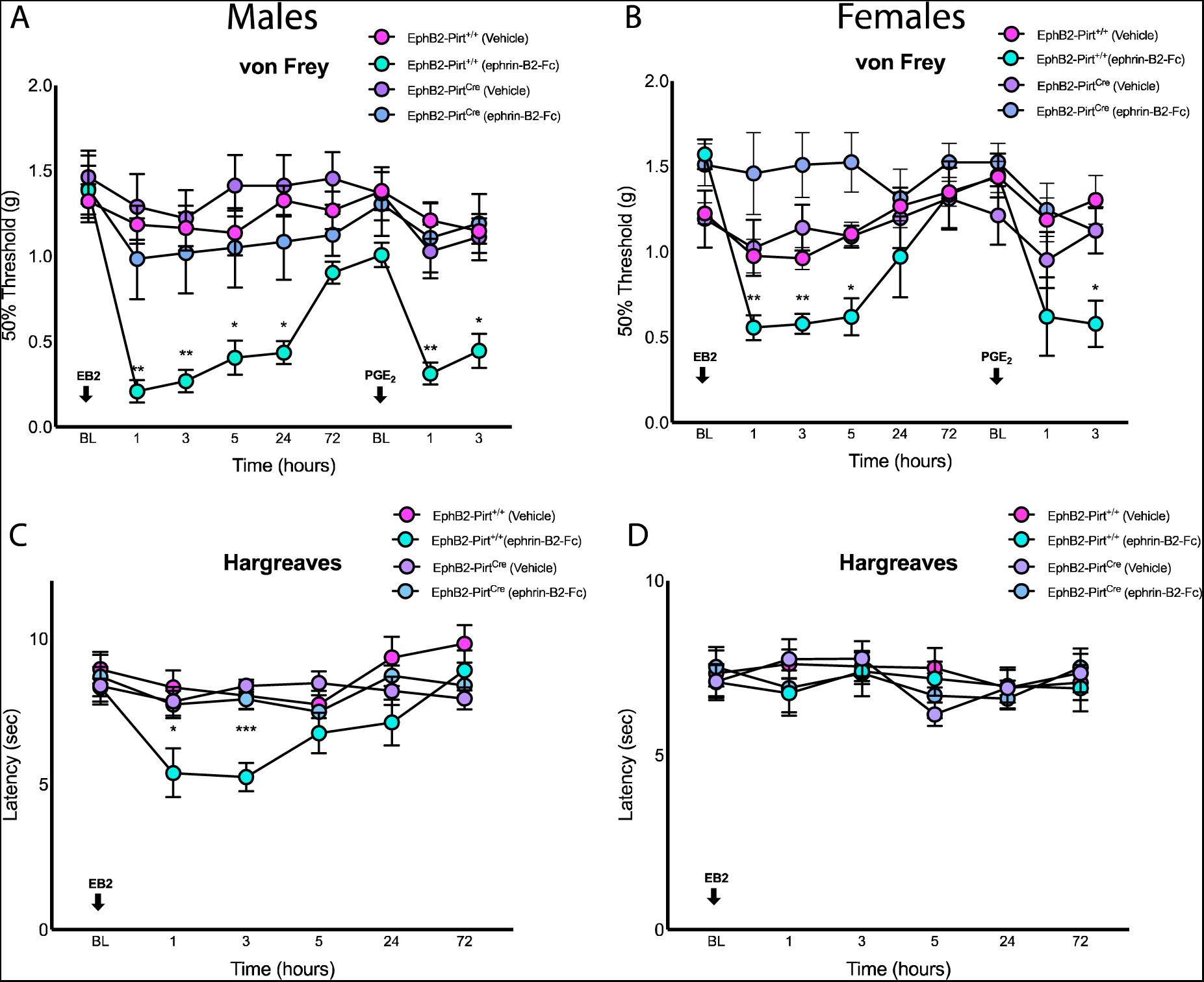
EphB2-Pirt^Cre^ mice show significantly reduced nociceptive behaviors in response to ephrin-B2 treatment. A, B, EphB2-Pirt^+/+^ and EphB2-Pirt^Cre^ mice were given an intraplantar injection of 100 ng of ephrin-B2-Fc and tested for mechanical hypersensitivity. After resolution of initial hypersensitivity, mice were given an injection of PGE_2_ to assess the presence of hyperalgesic priming. C, D, EphB2-Pirt^+/+^ and EphB2-Pirt^Cre^ mice were given a 100 ng injection of ephrin-B2-Fc intraplantarly and tested for thermal hyperalgesia. Males and females n = 6/group. EB2 = ephrin-B2-Fc. Repeated measures two-way ANOVA with Bonferroni’s *post hoc* test: *p < 0.05, **p < 0.01.

### Supradural ephrin-B2 treatment induces MNK1-dependent periorbital hypersensitivity and hyperalgesic priming in both sexes

Migraine headache is a neurological disorder characterized by intense head pain that is usually periodic and with episodes that can be caused by triggers [3; 20]. Mouse models of priming with stimuli applied to the cranial dura have been developed and have advanced our understanding of basic headache mechanisms [5; 7; 14; 15; 32; 33; 40]. We hypothesized ephrin-B2 would be sufficient to induce a migraine-like state in mice and that priming in response to ephrin-B2 would depend on MNK-eIF4E signaling. Male mice injected supradurally with a 100 ng dose of ephrin-B2 displayed periorbital mechanical hypersensitivity and only males that were previously treated with ephrin-B2 showed evidence of hyperalgesic priming when challenged with a subsequent dose of 100 ng PGE_2_ applied to the cranial dura (**Fig 5A**). Identical effects were seen in female mice (**Fig 5B**). To determine if these effects were driven through MNK-eIF4E signaling, MNK1^−/−^ mice were injected with 100 ng ephrin-B2. Male mice displayed significant acute periorbital mechanical hypersensitivity in both wild-type and MNK1^−/−^ groups, but only wild-type mice that received ephrin-B2 developed hyperalgesic priming (**Fig 5C**). Similar effects were seen in female mice, demonstrating that no sex differences are observed in this model with ephrin-B2 as the stimulus (**Fig 5D**).

**Fig 5:**
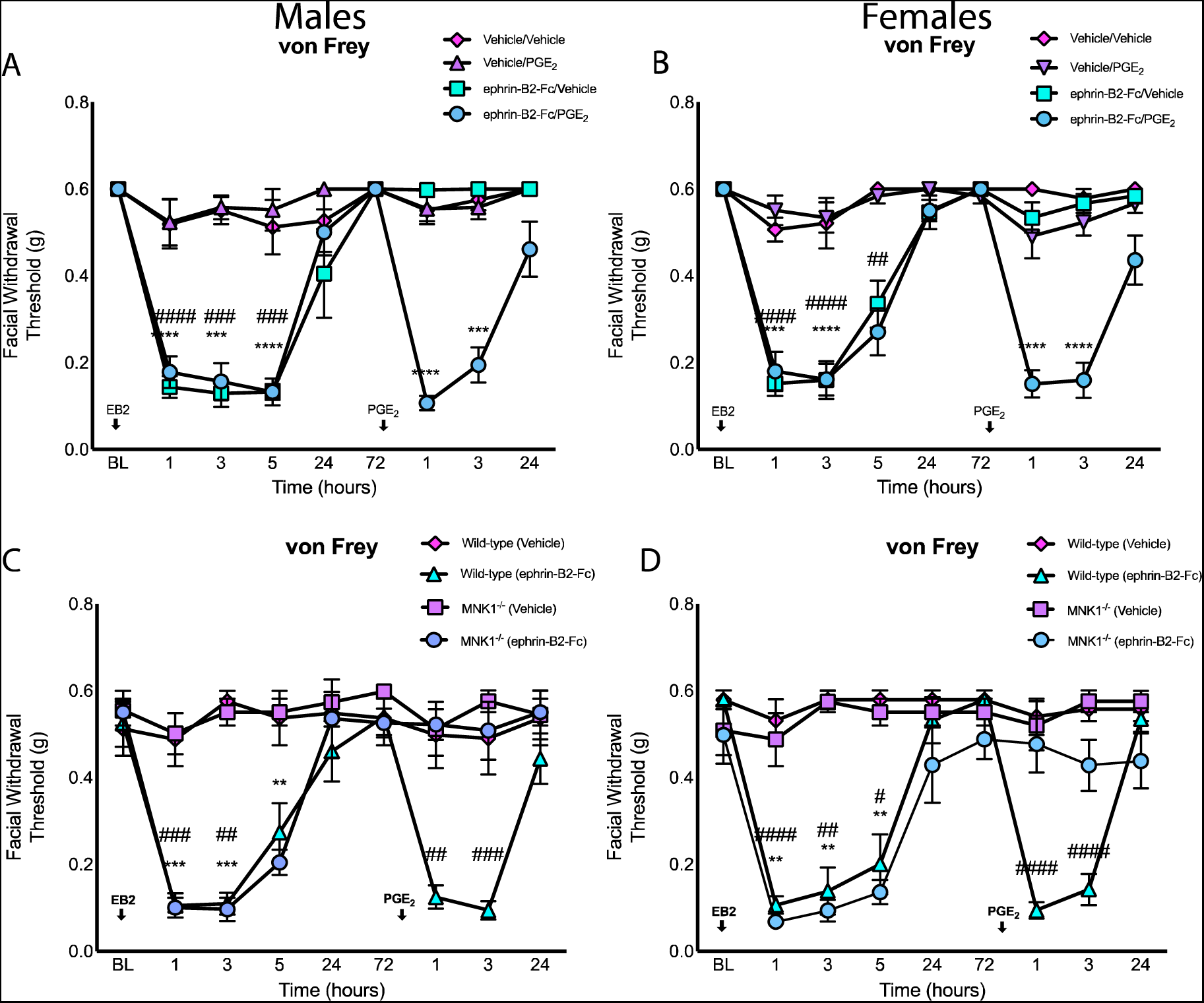
Supradural injection of ephrin-B2-Fc promotes peri-orbital mechanical hypersensitivity and priming that is MNK1-dependent. A, B, Naïve male (A) and female (B) mice were given a supradural injection of 100ng ephrin-B2-Fc or vehicle and periorbital mechanical hypersensitivity was assessed. After initial hypersensitivity resolved, mice were given a supradural injection of 100ng PGE_2_ or vehicle to assess the presence of hyperalgesic priming. Asterisks represent statistical significance of ephrinB2-Fc/PGE_2_ group versus vehicle/vehicle. # represents statistical significance of ephrinB2-Fc/vehicle versus vehicle/vehicle. Ephrin-B2/PGE_2_ (n = 8). Ephrin-B2-Fc/Vehicle and vehicle/vehicle (n = 6). Vehicle/PGE_2_ (n = 7). C, D, Wild-type and MNK1^−/−^ mice received a supradural injection of 100ng ephrin-B2-Fc or vehicle. Priming was determined through a dural injection of 100ng PGE_2_. Stars represent statistical significance of MNK1^−/−^ ephrin-B2-Fc versus Wild-type vehicle. Pound represents statistical significance of Wild-type ephrin-B2-Fc group versus Wild-type vehicle. Wild-type ephrin-B2-Fc (n = 7). All other groups (n = 6). EB2 = ephrin-B2-Fc. Repeated measures two-way ANOVA with Bonferroni’s *post hoc* test: *p < 0.05, **p < 0.01, ***p < 0.001, ****p < 0.0001.

### Ephrin-B2-Fc treatment sensitizes mouse DRG neurons to PGE_2_ in a MNK1- and EphB2-dependent fashion

We next sought to assess priming-like effects on DRG neurons in culture. Dissociated DRGs from both male and female mice were pre-treated with control IgG or ephrin-B2-Fc for 1 hour prior to stimulation with PGE_2_ (50 nM) and high K^+^ solution (50 mM) to examine changes in intracellular Ca^2+^ (**Fig 6A**). Pre-treatment with 0.35 µg ephrin-B2-Fc was sufficient to induce increased responses to PGE_2_ (representative traces in **Fig 6B**; analysis in **Fig 6C** (ephrin-B2-Fc treated neurons: X^2^ = 24.18, df = 1, p < 0.001) and **Fig 6D** (WT neurons: X^2^ = 22.91, df = 1, p < 0.0001)). Pre-treatment with ephrin-B2-Fc co-incubated with APV at 50 µM had no effect on responses to PGE_2_ demonstrating NMDA receptor engagement is not responsible for this priming effect (**Fig 6C**; APV treated neurons: X^2^ = 25.94, df = 1, p < 0.0001). On the other hand, this priming effect was not seen in DRG neurons from MNK1^−/−^ mice (**Fig 6D**; MNK1 neurons: X^2^ = 0.6228, df = 1, p = 0.4300). We determined if these *in vitro* results were driven through the EphB2 receptor by using ephrin-B2-Fc pre-treatment on DRG neurons from EphB2-Pirt^Cre^ and EphB2-Pirt^+/+^ mice. Only 7.69% of EphB2-Pirt^Cre^ responding to PGE_2_ treatment while 64.29% of neurons from mice with intact EphB2 responded to PGE_2_ (**Fig 6E**; EphB2-Pirt^+/+^ neurons: X^2^ = 14.96, df = 1, p < 0.0001; EphB2-Pirt^Cre^ neurons: X^2^ = 1.007, df = 1, p < 0.3156). We also assessed changes in latency to peak Ca^2+^ concentration times in PGE_2_-responding neurons. We found a significantly shortened latency to peak Ca^2+^ between wild-type vehicle and ephrin-B2-Fc pre-treated neurons, while no difference was found between MNK1^−/−^ vehicle and ephrin-B2-Fc pre-treated neurons. Additionally, we found that latency to peak in wild-type ephrin-B2-Fc pre-treated neurons was significantly shorter compared to both MNK1^−/−^ vehicle and ephrin-B2-Fc pre-treated groups (**Fig 6F**).

**Fig 6:**
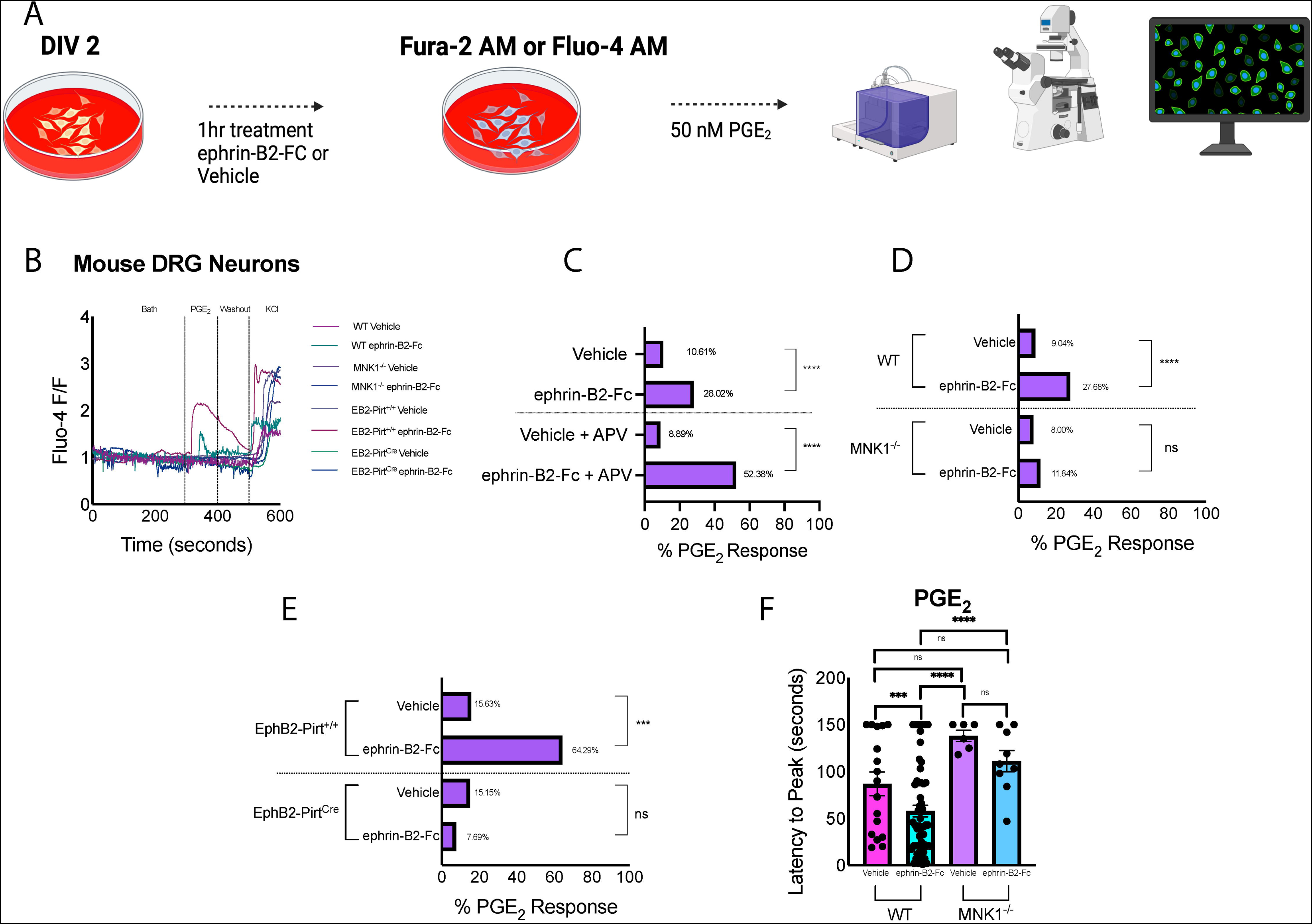
Ephrin-B2-Fc treatment sensitizes mouse DRG neurons in an EphB2 and MNK-dependent fashion. A, Diagram of the experimental timeline. B, Representative Ca^2+^ traces in response to PGE_2_ (50 nM) and K^+^ (50 mM). C, D, E, Total percent neuronal response rate to 50nM PGE_2_ treatment. Vehicle treated (n = 245). Ephrin-B2-Fc treated (n = 257). Vehicle + APV treated (n = 44). Ephrin-B2-Fc + APV treated (n = 126). WT vehicle treated (n = 188). WT ephrin-B2-FC treated (n = 224). MNK1^−/−^ vehicle (n = 75). MNK1^−/−^ ephrin-B2-Fc (n = 76). EphB2-Pirt^+/+^ vehicle (n = 32). EphB2-Pirt^+/+^ ephrin-B2-Fc (n = 28). EphB2-Pirt^Cre^ vehicle (n = 33). EphB2-Pirt^Cre^ ephrin-B2-Fc (n = 39). F, latency to peak was determined in ephrin-B2-Fc or vehicle treated cultures for both PGE_2_ (WT vehicle, WT ephrin-B2-Fc, MNK1^−/−^ vehicle, MNK1^−/−^ ephrin-B2-Fc, out of n = 17, 62, 6, and 9 responding neurons, respectively) and KCl positive neurons (WT vehicle, WT ephrin-B2-FC, MNK1^−/−^ vehicle, MNK1^−/−^ ephrin-B2-Fc, out of n = 188, 224, 75, and 76 responding neurons, respectively). Asterisks represent statistical significance between groups. WT = Wild-type; ns = not significant; EB2 = ephrin-B2-Fc. Bars represent standard error of mean. PGE_2_ % response, chi-squared analysis: ***p < 0.001, ****p < 0.0001. Ordinary two-way ANOVA with Bonferroni’s *post hoc* test: *p < 0.05, **p < 0.01, ***p < 0.001, ****p < 0.0001.

### Ephrin-B2 sensitizes human DRG neurons to PGE_2_, an effect blocked by eFT508 treatment

We next tested if ephrin-B2-Fc would cause sensitization of human DRG neurons in a MNK-dependent fashion. To do this we used an identical stimulation protocol to what we used in mice to assess sensitization of neurons (**Fig 7A**). Representative traces of Ca^2+^ responses evoked by PGE_2_ (50 nM) and 50 mM K^+^ treatment in each pre-treatment condition are shown in Fig 7B. The proportion of neurons responding to PGE2 were 3.15% in IgG Control pre-treated neurons, 34.19% with ephrin-B2-Fc treatment. No neurons responded to PGE_2_ with ephrin-B2-Fc and eFT508 (25 nM) co-incubation. (**Fig 7B**; IgG Control neurons compared to ephrin-B2-Fc: X^2^ = 48.05, df = 1, p < 0.0001. Latency to peak in response to PGE_2_ was significantly decreased in ephrin-B2-Fc pre-treated neurons (**Fig 7C**) and a change caused by eFT508 treatment could not be assessed because none of these neurons responded to PGE_2_.

**Fig 7:**
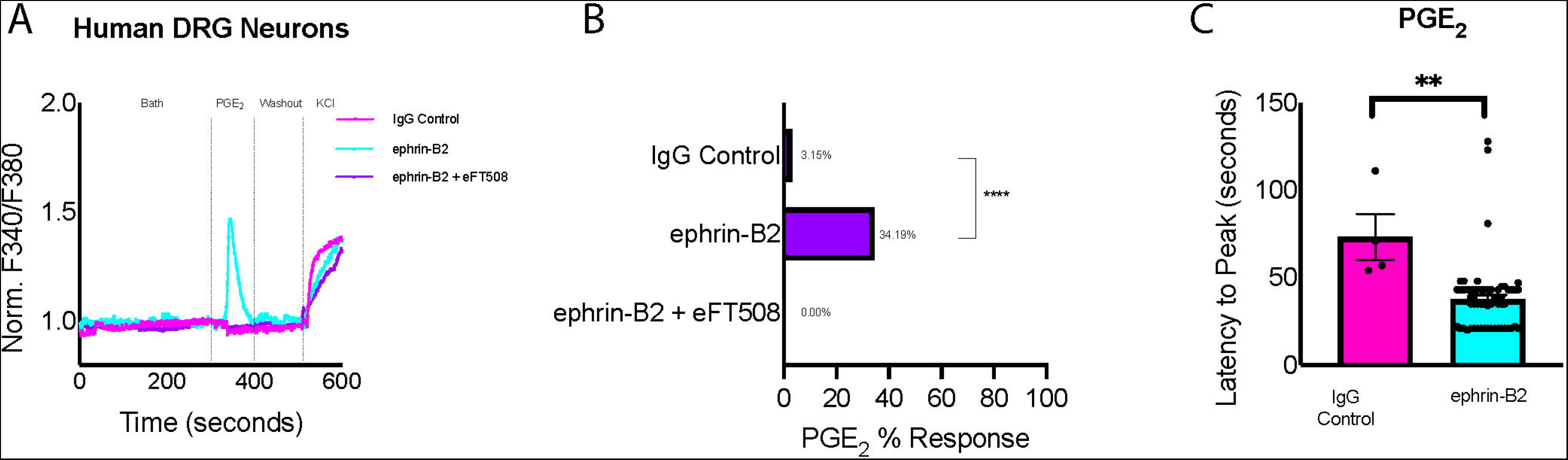
Ephrin-B2 treatment sensitizes human DRG neurons in a MNK-dependent fashion. Cultures of human dissociated DRG neurons were pre-treated with either ephrin-B2 (0.35 µg), IgG control, or ephrin-B2 and eFT508 (25 nM) prior to delivery of 50nM PGE_2_. A, Representative traces of pre-treated neurons to 50nM PGE_2_. B, Total response rate to 50nM PGE_2_ in dissociated cultures treated with IgG control or ephrin-B2, n = 4 (out of 134 total), 61 (out of 155 total), respectively. A total of 7 donors were cultured. C, latency to peak was determined in ephrin-B2 or vehicle treated cultures for PGE_2_ responsive neurons. Only neurons responsive to both PGE_2_ and K^+^ were assessed (IgG control and ephrin-B2, n = 4 and 61 responding neurons, respectively). Asterisks represent statistical significance between groups. Bars represent standard error of mean. Male (n = 3 independent cultures) and female (n = 4 independent cultures) groups were combined for analysis. PGE_2_ % response, Chi-squared analysis: ****p < 0.0001 PGE_2_ latency peak, Unpaired t test: **p < 0.01. K^+^ latency to peak, Ordinary one-way ANOVA with Bonferroni’s *post hoc* test: **p < 0.01, ****p < 0.0001. ns = not significant.

### Ephrin-B2 treatment increases eIF4E phosphorylation in human DRG neurons

Finally, we tested whether ephrin-B2-Fc treatment can induce MNK activation in human DRG neurons by measuring phosphorylation of eIF4E. Dissociated human DRG cultures were treated with IgG Control, ephrin-B2-Fc, or ephrin-B2-Fc and eFT508. We observed that ephrin-B2-Fc treatment induced a significant increase in eIF4E phosphorylation that was completely blocked by eFT508 (**Fig 8A** and **Fig 8B**). Consistent with the expression of EphB2 in most subsets of DRG neurons in humans [69], we observed that ephrin-B2-Fc treatment increased p-eIF4E in both large diameter neurons greater than 40 μm diameter (**Fig 8C**) as well as in small diameter neurons (<40μm; **Fig 8D**).

**Fig 8:**
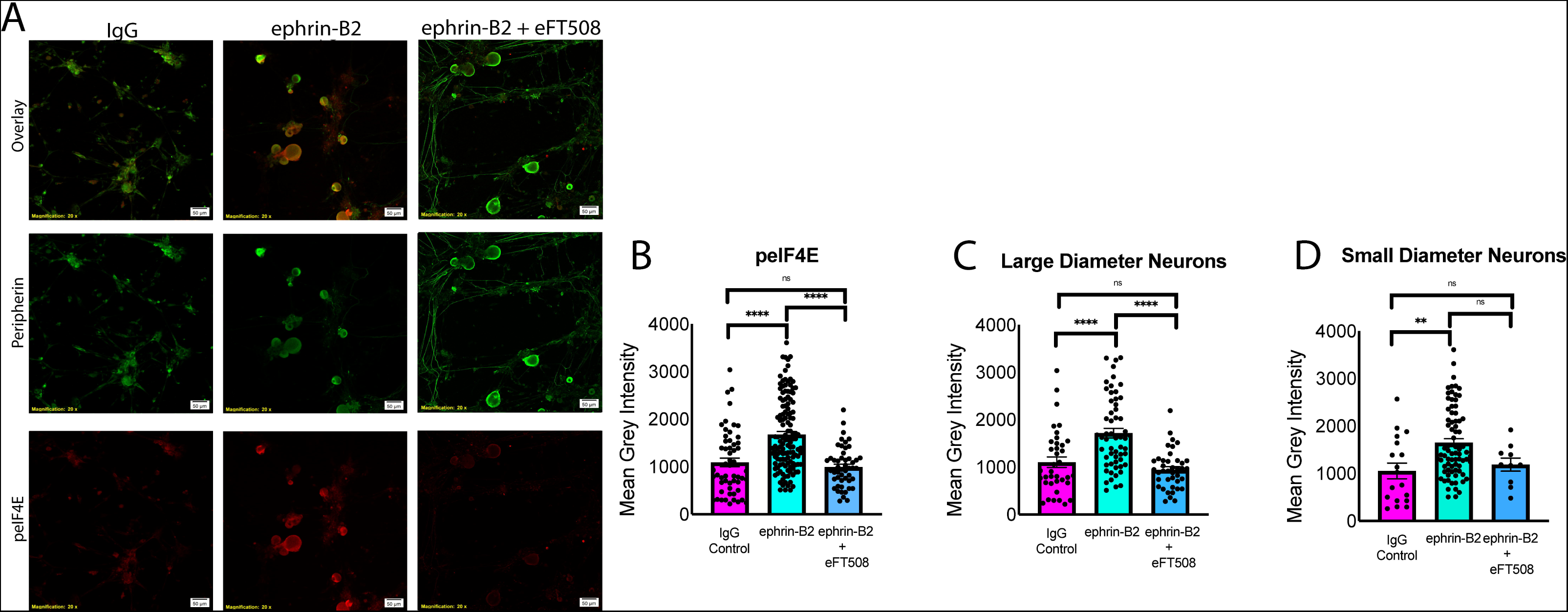
Ephrin-B2 causes increased eIF4E phosphorylation in human DRG neurons. Immunocytochemistry of dissociated human DRG cultures treated with either IgG control, ephrin-B2, or ephrin-B2 and eFT508 for 1 hour. A, Representative panel stained for Peripherin (green) or p-eIF4E (red). IgG control or ephrin-B2 treated at 0.35 μg/mL or ephrin-B2 and eFT508 (25 nM) treated neurons. B, Quantification of p-eIF4E expression. IgG control, ephrin-B2, ephrin-B2 and eFT508 treated neurons n = 55, 95, and 50, respectively. C, Quantification of large diameter neurons (≥40μm) p-eIF4E expression. IgG control, ephrin-B2, ephrin-B2 and eFT508 treated neurons n = 37, 57, and 40, respectively). D, Quantification of small diameter neurons (<40μm) p-eIF4E expression. IgG control, ephrin-B2, ephrin-B2 and eFT508 treated neurons n = 18, 77, 10, respectively). Stars represent statistical significance between groups. Bars represent standard error of mean. ns, not significant. Male (n = 3 independent cultures) and female (n = 2 independent cultures) groups were combined for analysis. Ordinary one-way ANOVA with Bonferroni’s *post hoc* test: **p < 0.01, ****p < 0.0001.

## Discussion

Our findings support the conclusion that ephrin-B2 acts directly on sensory neurons through the EphB2 receptor to cause mechanical sensitization, hyperalgesic priming and enhanced responses to PGE_2_ both *in vivo* (as reflected by hyperalgesic priming) and *in vitro*. All of these effects were mediated by MNK signaling, in both mice and humans providing important insight into the cellular signaling mechanisms driving these effects of ephrin-B2. These findings support the relevance of increased ephrin-B2 expression in painful diseases like osteo- and rheumatoid arthritis [8; 31] and pancreatic cancer [73] as a causative factor in driving pain through a direct action on DRG neurons that express EphB2.

Prior studies have shown ephrin-B2 can promote pain but these studies have either used male mice, or have not specified the sex of mice used in the studies [10; 24; 37; 66; 79]. Therefore, prior to this work it was unknown how male and female mice might differ in their response to ephrin-B2. Given pervasive sex differences in pain mechanisms [45; 46; 61] this represents an important gap in knowledge that needs to be addressed. We did observe a sex difference in heat hyperalgesia induced by ephrin-B2. Our findings in male mice were consistent with findings of peripherally delivered ephrin-B2 inducing heat hyperalgesia in male rodents [65]. We did not observe this heat hyperalgesia in female mice, and we do not know what the cause of the underlying difference might be. Nevertheless, we did not observe other sex differences, and we did not observe sex differences in effects on human DRG neurons. We surmise that effects in humans are likely to be consistent between men and women based on these findings.

A role of ephrin-B2 in migraine was recently suggested through brainstem signaling between TG afferents and second order neurons in the medullary dorsal horn [72]. Expression of EphB2 receptors in both mouse and human single cell sequencing data sets of the TG led us to hypothesize the involvement of ephrin-B2 in initiation of periorbital mechanical hypersensitivity and subsequent priming through signaling within the dura. Delivery of ephrin-B2 directly onto the dura was sufficient to induce these nociceptive behaviors related to migraine in both sexes. The priming effect was absent in MNK1^−/−^ but acute mechanical hypersensitivity was not attenuated, a finding that was not in line with observations of ephrin-B2 effects when given into the hindpaw. Previous studies have repeatedly shown that acute effects of many cytokines and growth factors are decreased in MNK1^−/−^ mice when the mediator is injected into the hindpaw [41; 48; 49]. The only previous study examining TG-associated signaling in MNK1^−/−^ mice found only small decreases to acute responses to interleukin-6 stimulation while stress-induced mechanical hypersensitivity was not changed in these mice [33]. Hyperalgesic priming, however, was completely absent in IL-6-treated or stressed mice [33], consistent with our findings in this study. While we do not know what the difference in the underlying acute mechanisms might be between the DRG and TG, and while we note that these neurons have important differences [13; 39; 42; 75], it is clear that hyperalgesic priming in both systems depends on MNK activity. One potential mechanistic difference is the hindpaw model measures primary mechanical hypersensitivity at the site of injection while the supradural model measures referred mechanical hypersensitivity in the periorbital area. The acute generation of mechanical hypersensitivity in the latter model likely requires central mechanisms while the creation of a primed state depends on plasticity in peripheral nociceptors and at central synapses [5; 6; 14; 32; 33]. MNK signaling may only influence the peripheral nervous system plasticity in the supradural model, consistent with previous observations [32; 33]. Another finding emerging from our work in the supradural migraine model is that PGE_2_ can be used as the stimulus to assess priming in this model as this had previously been untested. PGE_2_ has been shown to elicit headache symptoms in patients with migraine so this finding is consistent with clinical observations [2; 22; 47; 50].

Our study is not without limitations. First, as mentioned above we have not resolved the mechanisms that underlie either the sex difference in heat hyperalgesia between male and female mice, or the reason that acute, but not priming, effects of ephrin-B2 are differentially dependent on MNK1 signaling in the DRG system versus the TG. Second, while we show definitive evidence that ephrin-B2-medaited effects in mice depend on EphB2 expression in sensory neurons, we have not been able to test this in human DRG neurons due to a lack of available tools to inhibit EphB2. Finally, while our findings show that EphB2 plays a key role in the nociceptive effects induced by ephrin-B2, we cannot exclude a contribution of EphB1 in these effects as this receptor is also expressed in the mouse and human DRG [12; 37; 69; 78] and has previously been implicated in the peripherally-mediated effects of ephrin-B2 [37].

Our findings have important clinical implications. While previous research has focused on the very important role of ephrin-B-EphB signaling and its control of NMDAR activity in the dorsal horn of the spinal cord [10; 24; 30; 36; 66; 70; 72; 79], there is also evidence of a peripheral role of ephrin-B-EphB signaling in the peripheral nervous system in pain in animal models [37], and increased expression of ephrin-B2 in painful tissues in humans [8; 31; 73]. Given our observation of a loss of ephrin-B2-induced effects in a sensory neuron-specific EphB2 knockout mouse, one approach could be targeting of the EphB2 receptor in the periphery. Small molecule inhibitors of EphB1 have been developed and show efficacy in preclinical pain models suggesting a similar approach can be taken for EphB2 [1]. Another approach is inhibition of MNK activity, which would likely attenuate signaling via multiple pathways in addition to ephrin-B2-EphB2 signaling, such as cytokines and growth factors [33; 34; 41; 48; 49]. Given that these factors seem to upregulated in unison in painful diseases like rheumatoid arthritis [8], targeting MNK with near-clinical inhibitors of the kinase like eFT508 [21] may be a clinically effective way to control pain in rheumatoid arthritis.

Another clinically relevant aspect of our findings is the potential role of ephrin-B2-EphB2 signaling in the transition from acute to chronic pain. Preclinical models of this transition are controversial [56], in particular with respect to timing [51], but hyperalgesic priming models do mimic the change in nociceptive sensitivity that is often observed in patients even after an initial injury resolves [55; 58; 59]. To that end, it is notable that EphrinB2 treatment was capable of stimulating robust hyperalgesic priming and changes in dissociated DRG neuron physiological responses to both PGE_2_ and high K^+^ stimulation. The larger number of neurons responding the PGE_2_ likely reflects a change in the expression of GPCRs or signaling molecules downstream of these receptors. MNK activation, which controls translation of a subset of mRNAs important for inflammation and sensory neuron excitability [27; 28; 34; 41; 44; 48; 77], clearly plays a key role in this process. The change in the latency to peak for PGE_2_ and high K^+^ treatment caused by ephrin-B2 treatment likely reflects a change in the excitability state of DRG neurons. Again, this is consistent with observations in patients with chronic pain where nociceptors become hyperexcitable, even to the point of generating spontaneous activity [53] and this effect can also be reversed with MNK inhibition [34]. These findings are consistent with ephrin-B2-EphB2-MNK signaling being an excellent target for treatment of pain in diseases where chronic pain is linked to increased ephrin-B2 expression, like multiple forms of arthritis [8].

## Acknowledgments

The authors thank Anna Cervantes, Peter Horton and Geoffrey Funk for DRG recoveries from organ donors through the Southwest Transplant Alliance (STA). We also thank the STA for coordination of DRG recoveries. We thank the organ donors and their families for their gift. Funding for this study was from NIH grants NS115441 and NS111976 to MBD and TJP, NS104200 to GD, and NS065926 to TJP.

## Abbreviations

EphB, Erythropoietin-producing hepatocellular type-B receptor; EphB1, Erythropoietin-producing hepatocellular type-B receptor 1; EphB2, Erythropoietin-producing hepatocellular type-B receptor 2; ephrin-B, Eph receptor-interacting protein B; ephrin-B2; Eph receptor-interacting protein B2; MNK 1/2, Mitogen activated-protein kinase interacting kinases 1 and 2; eIF4E, eukaryotic initiation factor 4E; p-eIF4E, phosphorylated eukaryotic initiation factor 4E; DRG, Dorsal root ganglion; Pirt, Phosphoinositide interacting regulator of transient receptor potential channels; NMDAR, N-methyl-D-aspartate receptor; TG, Trigeminal ganglion; MAPK, mitogen-activated protein kinase; ERK, extracellular signal-regulated kinases; RNA, Ribonucleic acid; mRNA, Messenger ribonucleic acid; FRDU, 5-fluoro-2′-deoxyuridine uridine; NGF, Nerve growth factor; HBS, Hank’s balanced salt solution; PGE_2_, Prostaglandin E_2_; ICC; Immunocytochemistry; GPCRs, G-protein coupled receptors

